# Diverse and flexible behavioral strategies arise in recurrent neural networks trained on multisensory decision making

**DOI:** 10.1101/2023.10.28.564511

**Authors:** Thomas Wierda, Shirin Dora, Cyriel M. A. Pennartz, Jorge F. Mejias

## Abstract

Behavioral variability across individuals leads to substantial performance differences during cognitive tasks, although its neuronal origin and mechanisms remain elusive. Here we use recurrent neural networks trained on a multisensory decision-making task to investigate inter-subject behavioral variability. By uniquely characterizing each network with a random synaptic-weights initialization, we observed a large variability in the level of accuracy, bias and decision speed across these networks, mimicking experimental observations in mice. Performance was generally improved when networks integrated multiple sensory modalities. Additionally, individual neurons developed modality-, choice- or mixed-selectivity, these preferences were different for excitatory and inhibitory neurons, and the concrete composition of each network reflected its preferred behavioral strategy: fast networks contained more choice- and mixed-selective units, while accurate networks had relatively less choice-selective units. External modulatory signals shifted the preferred behavioral strategies of networks, suggesting an explanation for the recently observed within-session strategy alternations in mice.

## Introduction

Individuals usually display a high level of variability in their cognitive abilities and goal-directed behavior^1,2^, often exceeding what would be expected even from genetically identical subjects^3–5^. When performing cognitive tasks such as decision making^6–9^ or multisensory integration^10–15^, individuals have often been shown to develop separate choice strategies^16–20^. In the case of mice performing decision-making tasks, Pittaras and colleagues observed a broad distribution of behavioral strategies depending on the risk-aversiveness of the animals^17^. Rats performing context-dependent decision-making tasks display individually different behavioral strategies, even when their performance levels were similar^18^. Behavioral strategies are, furthermore, not an invariant quality for a given animal, as mice have been shown to alter their decision strategy within the same session^19,20^. While the mechanisms underlying different behavioral strategies are still unclear, there is evidence of behavioral strategies being reflected in neurobiological features^17^, suggesting a connection between behavioral variability and neural and circuit heterogeneity, which is already known to play a beneficial role in neural processing and computations^21–25^.

In spite of its prevalence and impact, this inter-subject behavioral variability is still treated in most studies as noise, and averaged out across subjects in behavioral and neuroscience research on both human and non-human animal studies^1,2^. In the absence of systematic behavioral datasets or standardized protocols to control for behavioral variability in rodents, computational models offer a practical way to explore the impact of behavioral variability in experiments^18^. Models of recurrent neural networks (RNNs) are particularly well suited for this task, since they can be easily trained to perform simple perceptual, cognitive and behavioral tasks mimicking the ones used in actual experiments^26–32^. Unfortunately, most computational studies also average over different RNN realizations to present cohesive results, and therefore the potential of RNNs to uncover the origin and role of inter-subject behavioral variability has been largely untapped.

In this work, we trained a large number of RNNs on a simplified version of a classical multisensory stimulus discrimination task^11,33^ and explored the emergence of inter-subject variability in behavioral strategies within these networks. Each network was initialized with a unique set of synaptic weights before training, to reflect the natural variability present across subjects. The RNNs also included excitatory and inhibitory neurons (with a ratio of 4-to-1) to account for Dale’s law and enhance their biologically plausibility^28^. We found that training these initially heterogeneous RNNs under the same protocol led to a large variability in terms of reaction time, choice bias and accuracy of decisions, as reflected by their individual psychometric and chronometric curves. In addition, neurons within each network showed heterogeneous responses and could be classified as modality-selective, choice-selective, mixed-selective (i.e. modality and choice), hyper-selective, or silent^34^. There was a clear correlation between a network’s behavioral strategy and its neural composition: fast-deciding networks differed from slow ones in the proportion of choice-selective, mixed-selective and silent neurons, and accurate networks differed from inaccurate ones in the proportion of choice-selective units. The structure of these functional signatures was also different for excitatory and inhibitory neurons, suggesting differentiated roles between both classes^35^. Finally, we found that external modulatory currents were able to shift the preferred strategy of a given network, suggesting that neuromodulatory signals (such as acetylcholine or noradrenaline) or gating signals from other brain areas (such as thalamic nuclei) could be behind the within-session alternations in strategies observed in mice^19^.

## Results

We trained 100 RNNs (Fig. 1a) on a simplified version of a standard multisensory integration task^11,33^, in which the animal had to decide whether the rate of a series of sensory pulses was higher or lower than a known reference level (Fig. 1b). The pulses corresponded to flashes for visual trials, clicks for auditory ones, and combined flashes-clicks for audiovisual ones. In our model, each network consisted of 150 rate-based neurons, of which 80% were excitatory and 20% inhibitory^28^. Each network received two streams of input (one corresponding to visual input and the other to auditory input), and two output variables were produced from the excitatory population, coding for the two choices the network could make (high vs low rate). Sensory input was modeled as noisy step currents (Fig. 1c), whose amplitude represented visual and/or auditory pulse frequencies between 9 Hz and 16 Hz, with 12.5 Hz corresponding to the threshold between both options. On unimodal trials, the network received information for one modality, whereas on multimodal trials the network received congruent information about the frequency in both modalities (Fig. 1c). The network made a categorical decision by raising the correct output variable while keeping the other low (Fig. 1d). The stimulus was presented for 1 second after a fixation period of 100 ms in which the network had to keep both output variables low. A choice was recorded when the output variables were separated by more than 0.2 (given in arbitrary units for neural activity, with alternative values explored in Fig. S1), and a trial was considered invalid if a choice was registered before stimulus onset. On average, networks showed a high fraction (> 0.9) of valid trials after training on all frequencies, with a small dip in the number of valid trials around the threshold frequency (Fig. 1e). After training, we generated 8192 trials per model with equal probability across all conditions. Networks were generally able to perform the task well, with a decrease in performance for difficult trials presenting a frequency close to the threshold frequency (Fig. 1f, see Methods for more details).

**Figure 1.**
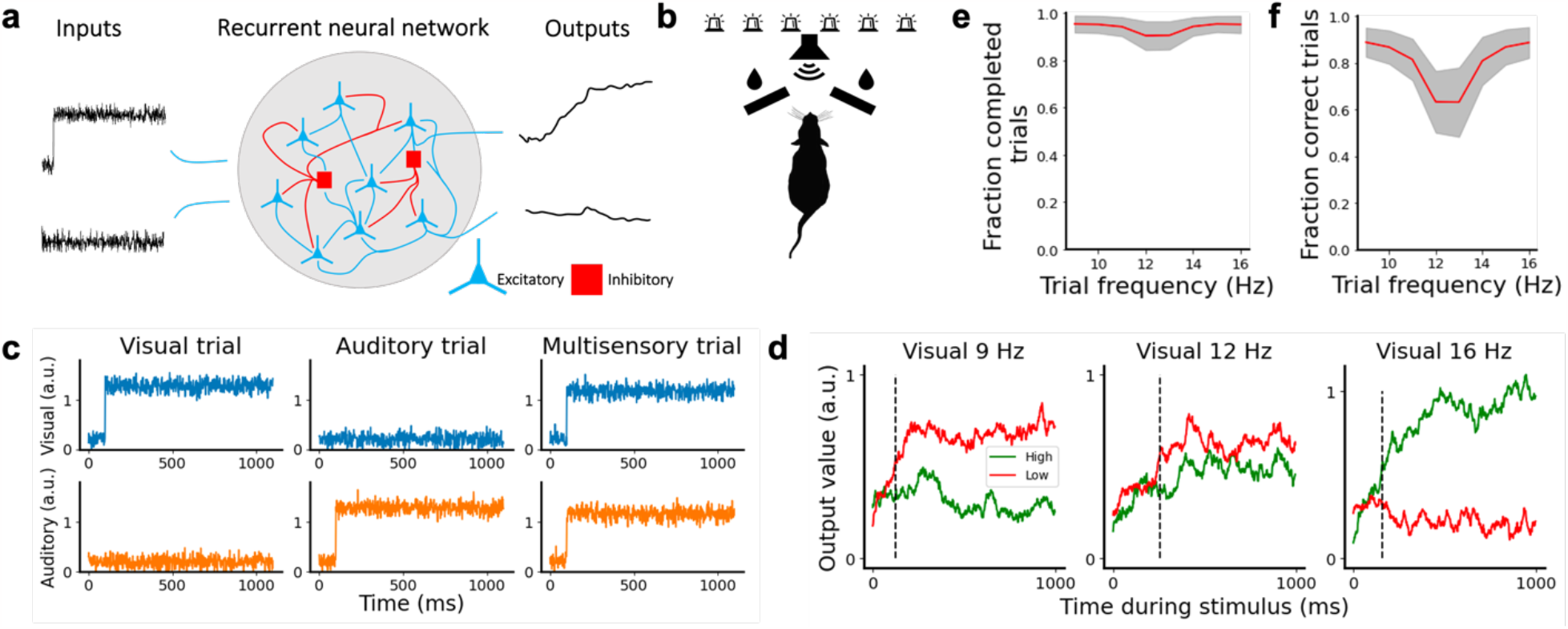
Recurrent neural network to model multisensory integration. **a** We considered 100 RNNs, each of them having 150 neurons (80% excitatory neurons and 20% inhibitory neurons), which received two streams of input and were trained to decide if the level of the input was above or below a threshold. The networks reported a choice by elevating the corresponding output variable. **b** Experimental task proposed by Raposo et al. and modeled in this work. In the original task, rats received visual flashes and/or auditory clicks and had to report whether the frequency was higher of lower than an experimenter-imposed threshold frequency. **c** Examples of input received by the model on visual (left), auditory (center) and multimodal (right) trials, via the visual (top) or auditory stream (bottom). **d** Examples of a network’s output variables throughout the trial on easy low (left), difficult (center), and easy high (right) trials. Dashed lines indicate the decision time of the trial. **e** Mean fraction of valid trials a network had per trial frequency, with the grey band indicating the s.e.m. across networks. **f** Mean fraction of trials with a correct decision for different trial frequencies.

### Networks display a significant variability in task performance

To characterize the behavioral output of each network, we calculated its psychometric curves (i.e. fraction of times the network chose ‘high’ as output vs input frequency) and chronometric curves (time to reach a decision vs input frequency) after training. For psychometric curves, only trials in which a decision was made during stimulus presentation were included. The chronometric curves were calculated based on trials resulting in a correct choice only. On average, networks showed improvement in their performance (Fig. 2a) and needed less time to form a decision (Fig. 2b) on multimodal trials as compared to unimodal trials, replicating experimental findings^11,33^. Notice that all networks were trained with the same protocol, and their behavioral differences are a consequence of the unique initial conditions (i.e. randomized weights) of each network before training, reflecting innate or pre-training variability across subjects. Furthermore, the 100 networks showed great variability in both their psychometric functions (Fig. 2c) and reaction times (Fig. 2d). This indicates that different networks could potentially adopt different behavioral strategies to solve the task, such as a fast-but-inaccurate strategy, a slow-but-accurate strategy, a preferentially-choose-low strategy, etc. We then sought to explore whether this was indeed the case, and whether structural differences were underlying the different strategies adopted by networks.

**Figure 2.**
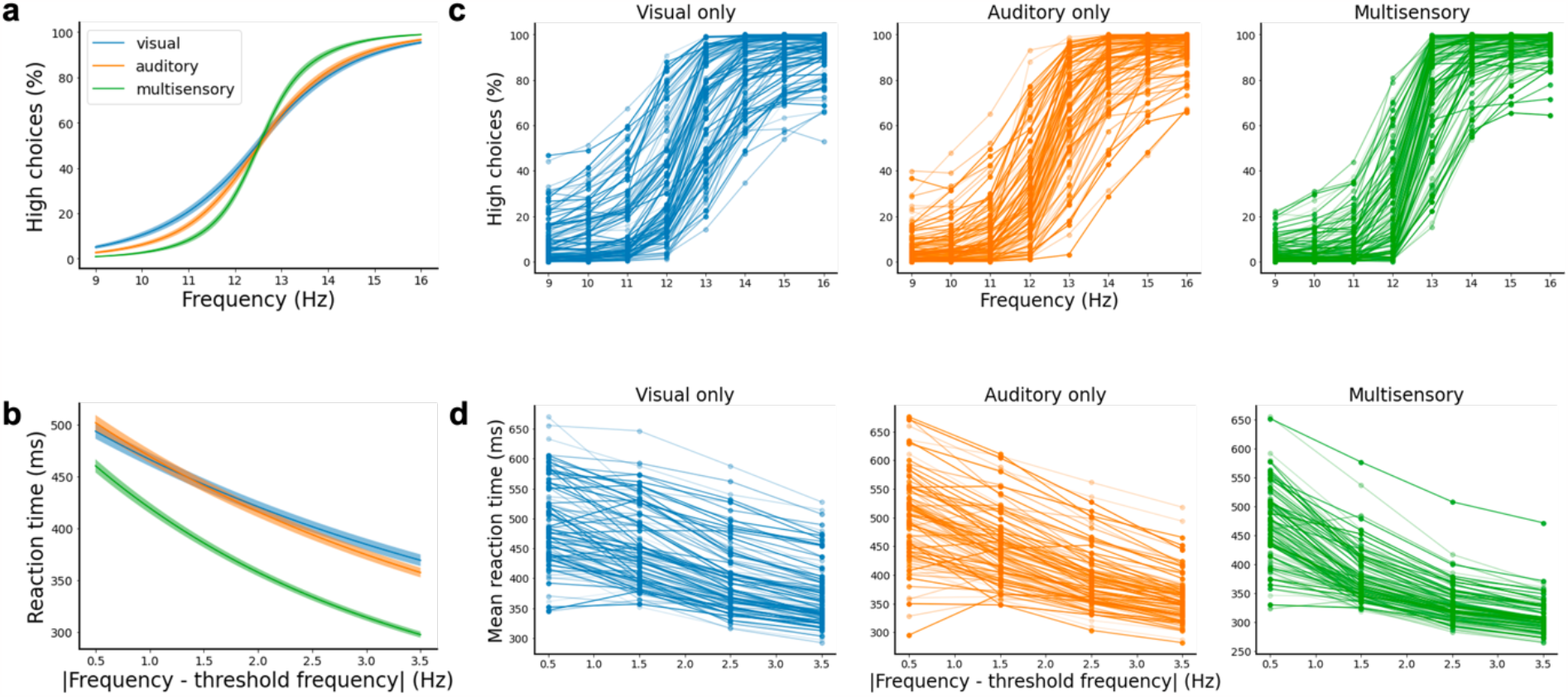
Networks show variability in task performance. **a** Psychometric functions averaged over all networks for visual (blue), auditory (orange) and multisensory (green) trials, where an improvement of multisensory over unisensory trials can be observed. Shaded bands represent the s.e.m. across networks. **b** Chronometric functions averaged over all networks on visual (blue), auditory (orange) and multisensory (green) trials; lower reaction times are observed for multisensory compared to unisensory trials. **c** Psychometric functions for individual networks, displaying a large variability across them. **d** Chronometric functions for individual networks, displaying a large variability across them.

### Multisensory trials lead to heterogeneous, but faster and more accurate responses

We used the psychometric and chronometric functions of the networks to map network performance to several behavioral metrics. For example, the slope parameter of the psychometric curve gives an indication of how well-defined the decision threshold of the network is (Fig 3a, left panel). A large slope value indicates that the network can confidently classify each input into one of the two outputs, whereas a small slope suggest that the network makes more mistakes in the classification of inputs of intermediate frequencies. We used this as a measure of network accuracy. The frequency offset parameter of the psychometric curve gives an indication of the frequency at which the network switches from low to high choices. We therefore used this as a measure of choice bias. To assess the decision speed of a network, we used the mean reaction time of all the network’s correct choices (Fig. 3a, middle panel). The combination of accuracy and choice bias (Fig. 3a, right panel) with the mean reaction time constitutes our full set of metrics to characterize a network’s behavioral strategy.

**Figure 3.**
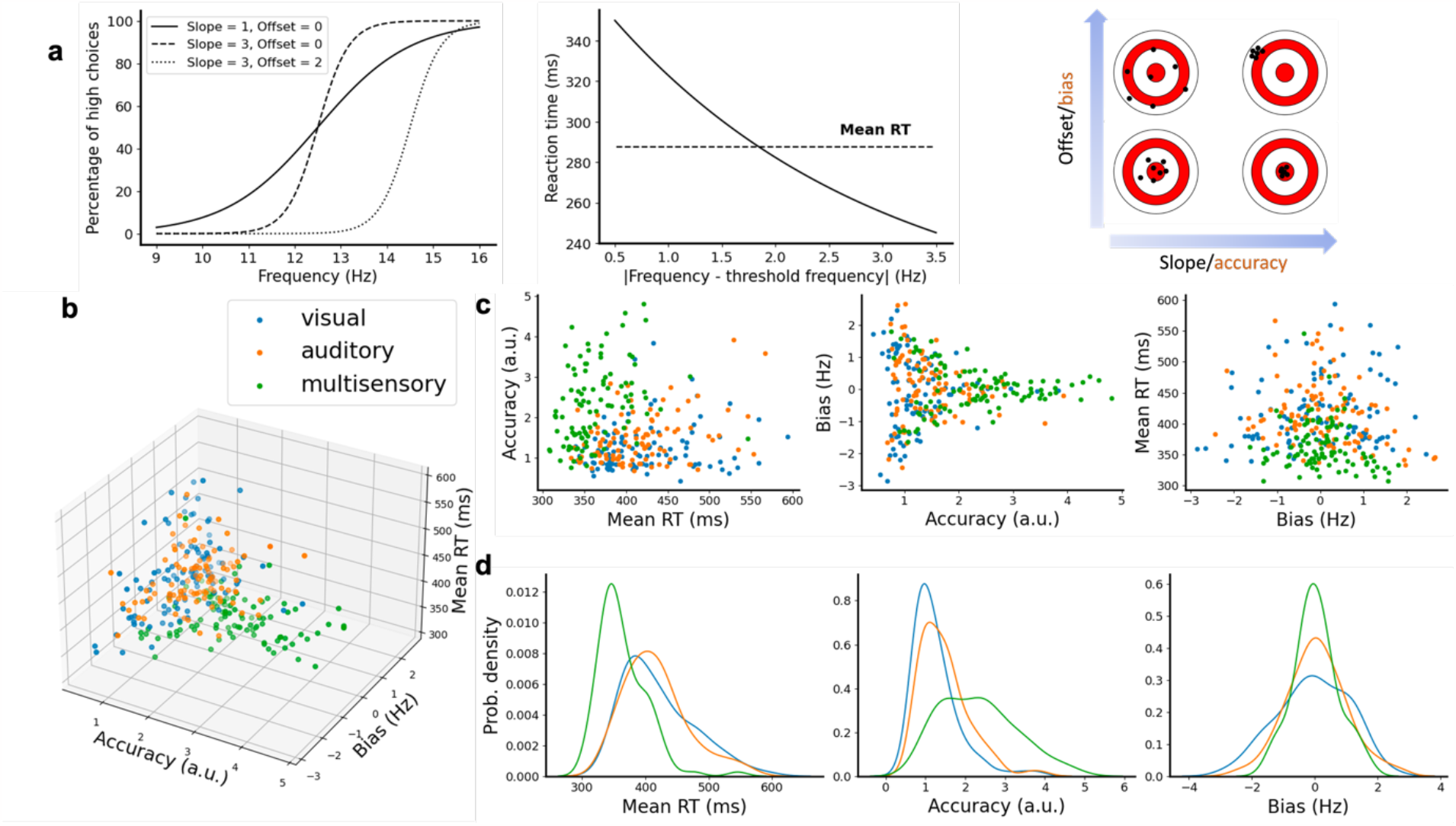
Multisensory trials lead to faster and more accurate network responses. **a** Sigmoidal (left panel) and exponential curves (middle panel) used to fit each network’s psychometric and chronometric data. The slope and frequency offset of the sigmoidal curves were used as fitting parameters and to estimate, respectively, the accuracy and choice bias of each network (right panel). The decision speed was estimated by computing the mean value of the reaction times for correct trials. **b** A 3D scatter plot reveals significant behavioral variability across all 100 networks and three conditions (visual, auditory and multisensory trials). **c** 2D projections of the 3D scatter plot for different pair of metrics. **d** The distribution of reaction times, accuracy and choice bias for visual, auditory and multisensory trials.

When mapped over the different trial modalities (visual, auditory and multisensory) we observed that networks showed a high variability in all metrics across the different modalities (Fig 3b). We also observed that some networks had a low accuracy with a high reaction time, whereas others had a high accuracy with a low reaction time. High accuracy with low reaction time was predominantly achieved on multisensory trials (green dots) rather than unimodal trials (orange/blue dots). We observed a positive correlation between the accuracy and the time it took to make a decision (Fig. S2), indicating a speed-accuracy trade-off^36–38^.

To explore these relationships more precisely, 2D-projections of the 3D scatter plot were scrutinized, comparing pairs of metrics (Fig. 3c). Low mean reaction times and high accuracy decisions were mostly achieved in multisensory integration trials, while unisensory trials led to slower and less accurate decisions (Fig. 3c, left). Comparing the accuracy with the choice bias (Fig. 3c, center) showed that the multisensory trials resulted in a wider range of accuracy values than those for unisensory trials, and that the more accurate a network was, the less biased they became. Comparing the bias with the mean reaction time (Fig. 3c, right) again demonstrated that networks generally decided faster on multisensory trials, but not at the cost of choice bias. To further quantify these 2D-projections we calculated the distributions of the different metrics (Fig. 3d). In multisensory trials the distribution of the mean reaction time was skewed more to the left than for either type of unimodal trial, with no major differences between unimodal trials (Fig. 3d, left). The distribution of accuracy values is wider for multisensory trials as compared to unimodal trials, with approximately the same minimum value but a larger maximum value (Fig. 3d, center). The distribution plot of choice bias showed that the networks achieve the lowest bias mostly on multisensory trials (Fig 3d, right). Fitting a more extended sigmoidal curve with lapse parameters, which account for shifts in percentage values of psychometric curves, gave similar results (Fig. S3 and Table S1).

### Network units show heterogeneous responses and vary between strategies

Upon examining the dynamics of neurons within the networks, we observed that they showed heterogeneous responses to different unimodal trial conditions. We classified neurons based on their firing rate activity on correct unimodal trials and distinguished five different types of selectivity: (1) modality selectivity, (2) choice selectivity, (3) mixed selectivity (i.e. a combination of modality and choice selectivity^34^), (4) silent or very-low-activity neurons, and (5) hyper-selectivity (Fig. 4a). To assess whether networks showing different strategies differed in their underlying physiological response properties and functional structure, we categorized the networks into groups based on their overall accuracy and mean reaction time irrespective of modality. A dichotomy between fast (N=60) versus slow (N=40) and accurate (N=39) versus inaccurate (N=61) networks was adopted based on the average of the metrics across all networks. Next, the number of selective units of the different types was compared between the groups, where differences could suggest that these units play a role in the formation of a network’s strategy. Fast networks did not differ significantly from slow networks in the number of modality- and hyper-selective units but did significantly differ in their number of choice-selective (p < 0.05, permutation test with Holm-Bonferroni correction unless specified otherwise), mixed-selective (p < 0.001) and silent neurons (p < 0.001)(Fig. 4b). More precisely, fast networks had more choice- and mixed selective neurons, but less silent neurons as compared to slow networks. There were less significant differences between accurate and inaccurate networks (Fig. 4c), with the only exception being that accurate networks had significantly less choice-selective neurons than inaccurate ones (p < 0.01, see Table S2).

**Figure 4.**
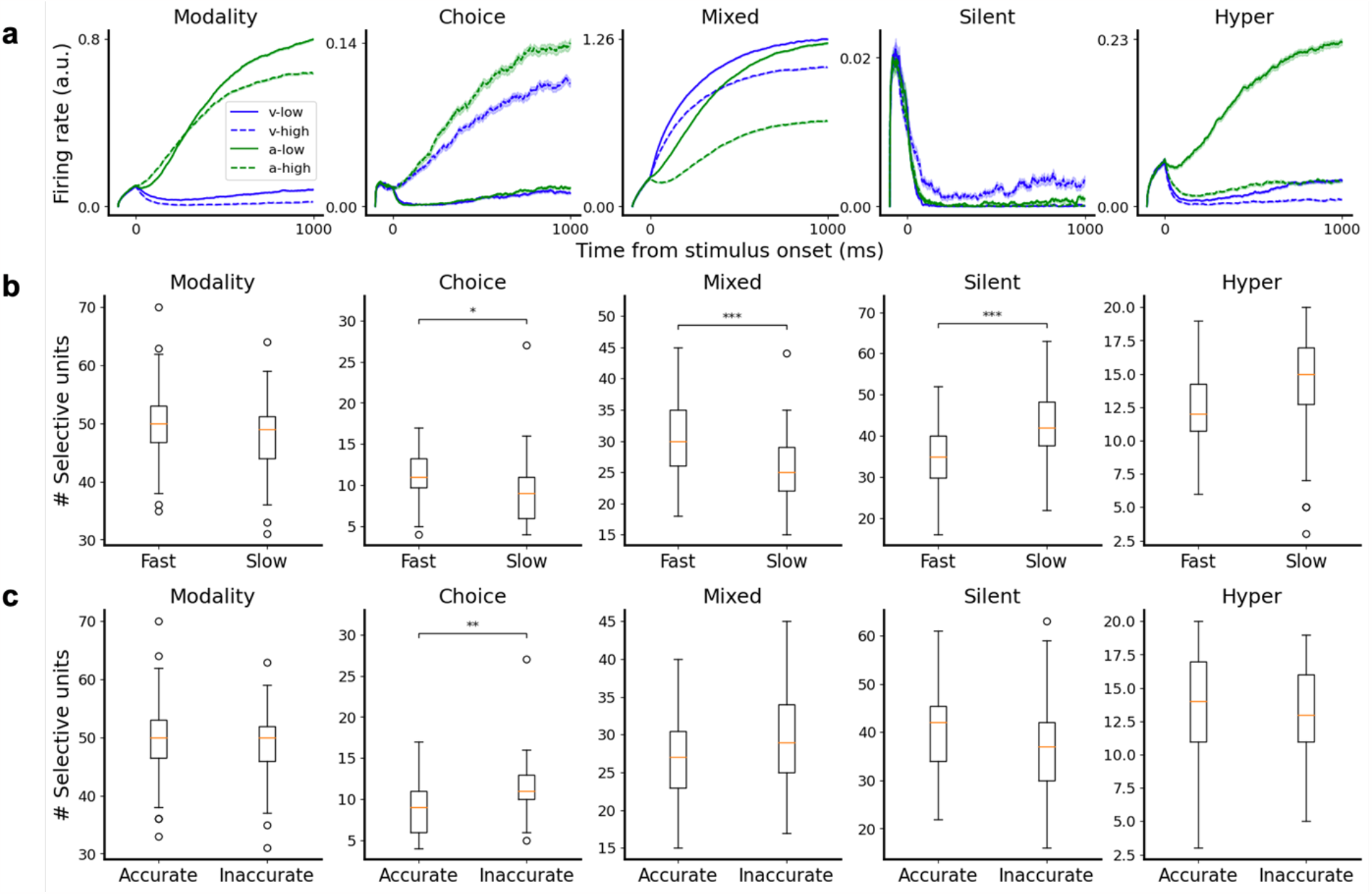
Network units show heterogeneous responses and vary between strategies. **a** Example of network units showing modality selectivity, choice selectivity, mixed selectivity, silence during the stimulus and hyper-selectivity. **b** Fast networks had significantly more choice- and mixed selective units and less silent neurons compared to slow networks. **c** Accurate networks had significantly less choice selective neurons compared to inaccurate networks. Significance: * p < 0.05, ** p < 0.01, *** p < 0.001 two-tailed permutation test with n=100,000 resamples and Holm-Bonferroni correction.

The absolute number of selective units was furthermore associated with reaction times and accuracy of the networks (Table S3): the number of choice- and mixed-selective units was negatively associated with the accuracy of the networks, whereas the number of silent neurons was positively associated with it. Having more modality-, choice- or mixed-selective units decreased a network’s reaction time, whereas having more silent and hyper selective neurons led to increased reaction times and thus slower networks.

### Inhibitory and excitatory populations differ in selectivity

On a population level, we observed that the fraction of mixed-selective neurons in the excitatory populations was significantly larger than the fraction in the inhibitory population (p < 0.001) (Fig 5a, Table S4). Inhibitory populations also displayed a significantly larger fraction of hyper-selective neurons than excitatory populations (p < 0.05). To further investigate whether the differences in strategies between networks could be attributed to differences between inhibitory and excitatory populations, we performed multiple comparisons both within and between groups. Within-group comparisons revealed that fast networks had on average a higher fraction of choice- and mixed-selective neurons in their excitatory population compared to their inhibitory population (Fig. 5b, Table S5) and that inaccurate networks had a lower fraction of mixed-selective neurons in their inhibitory population as compared to their excitatory population. The between-group comparisons revealed that the excitatory population of fast networks differed from the ones in slow networks in their fraction of choice-selective (p < 0.01), mixed-selective (p < 0.001) and silent neurons (p < 0.001). Accurate networks displayed a lower fraction of choice-selective excitatory neurons than inaccurate networks (Fig. 5c, p < 0.01, Tables S6), indicating that having a large fraction of choice-selective neurons does not necessarily lead to better performance. No significant differences were found between the groups in the inhibitory populations.

**Figure 5.**
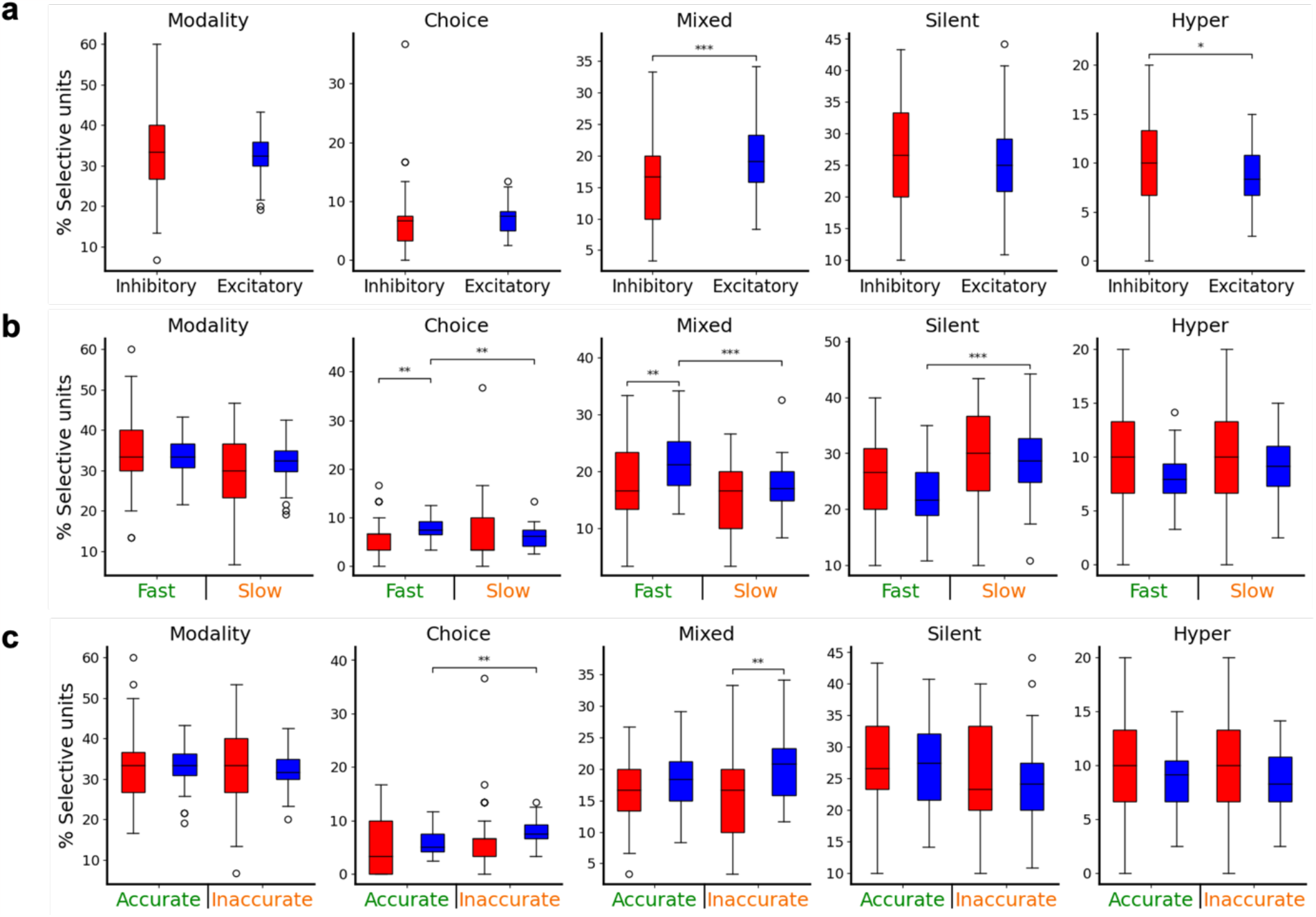
Inhibitory and excitatory populations differ in selectivity. **a** Comparison between the number of excitatory and inhibitory neurons found accross all five selectivity types. **b** Fraction of fast vs slow networks for both excitatory and inhibitory neurons and across all selectivity types. **c** Fraction of accurate vs inaccurate networks for both excitatory in inhibitory neurons and across all selectivity types. Significance: * p < 0.05, ** p < 0.01, *** p < 0.001 two-tailed permutation test with n=100,000 resamples and Holm-Bonferroni correction.

### Modulatory currents alter the adopted behavioral strategies

To explore whether our model was able to explain sudden changes in a subject’s behavioral strategy, as experimentally reported^19,20^, we studied the effect of adding an extra modulatory (constant) current to the network (Fig. 6a), which is aimed to reflect internal gating or neuromodulation. Across eight different experiments, eight different groups of neurons received this modulatory current throughout the entire trial duration. The eight groups that received the current were: all five different types of selective units, modulated one at a time (1-5), all neurons (6), inhibitory neurons only (7) and excitatory neurons only (8). The application of a modulatory current could alter the accuracy, bias and decision speed of the network, or in the terminology of Ashwood et al.^19^, lead to a network becoming more engaged, disengaged, biased or exhibiting changes in reaction times (Fig. 6b). Reaction times decreased when the modulatory current targeted modality-selective, choice-selective, mixed-selective, excitatory, or all neurons, while they increased with input targeting inhibitory neurons. Targeting silent or hyper-selective neurons did not affect reaction times (Fig. 6c, Table S7). On the other hand, accuracy decreased when the modulatory input targeted modality-selective, silent, hyper-selective, excitatory or all neurons, increased when the input targeted inhibitory or choice-selective neurons, and remained constant when targeting mixed-selectivity neurons. This leds to two interesting traits: (i) speed-accuracy trade-offs were observed when the modulatory signal targeted either modality-selective neurons, excitatory, inhibitory, or all neurons, and (ii) modulating choice-selective neurons led to improvements in both accuracy and decision speed. The modulatory current did not cause any significant shift in the bias of the models. Although on an individual network level we found changes in bias levels (Fig. 6d, left), the direction of the shift differed a lot between the networks resulting in no significant average shift to either direction. Similar results were found when considering other criteria for neural selectivity, such as one based on ROC curves^33,35^ (Fig. S4, Table S8).

**Figure 6.**
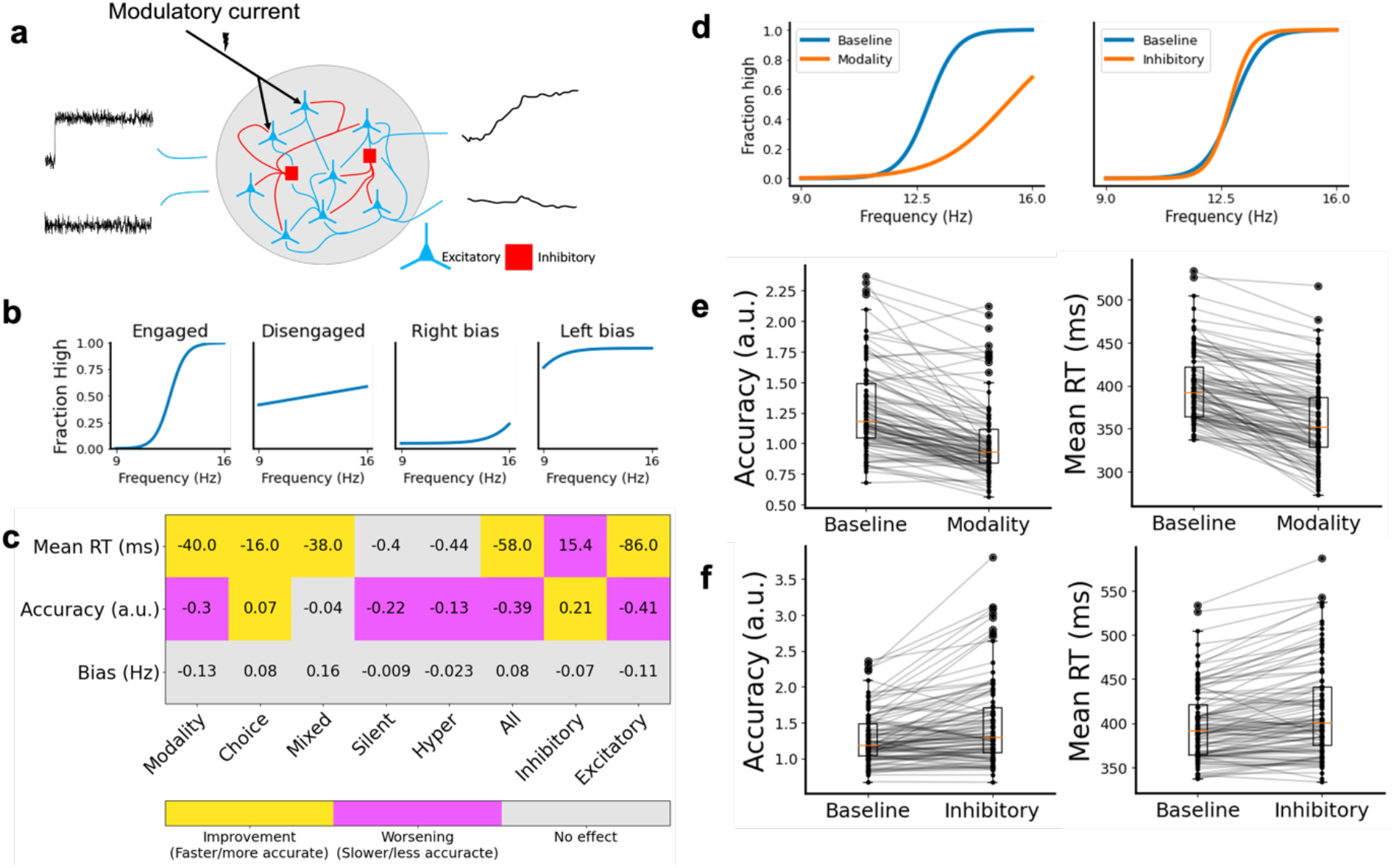
Modulatory currents can influence individual network strategies. **a** We applied a continuous modulatory current to different neuron subsets within a network to assess whether external modulation could influence individual behavioral strategies. **b** Potential effects of external modulation included networks becoming more engaged, disengaged, biased, and influence their reaction times. **c** Behavioral changes (as changes in reaction times, accuracy and bias) observed when the modulatory input targeted different groups (modality-selective, choice-selective, etc). Significant improvements (i.e. faster or more accurate decisions) are displayed in yellow blocks, worsening in magenta, and non-significant changes in grey. **d** Example of an individual network’s change in psychometric curves on the application of a modulatory current to its modality-selective (left) and its inhibitory (right) neurons. **e** Effect of modulatory input targeting modality-selective neurons in the accuracy (left) and reaction times (right) across all networks. **f** Effect of modulatory input targeting all inhibitory neurons in the accuracy (left) and reaction time (right) across all networks. The effects in panels e and f are opposite to each other, and may serve to rebalance speed-accuracy trade-offs in the network’s behavior.

An example network, shown in Fig. 6d, illustrates the opposing example of the effects that a modulatory current has on the network’s accuracy when targeting a group. In the first example, which considers an input targeting modality-selective neurons, the modulatory current lowers the accuracy (and introduces a right-shift bias) with respect to the baseline or control case. In the second example, in which the input targets all inhibitory neurons, the current increases the accuracy of the network. Changes in accuracy and reaction times across all networks for these two modulation examples are shown in Figs. 6e and 6f. These examples demonstrate that externally modulating modality-selective neurons or inhibitory neurons lead to speed-accuracy trade-offs of opposing signs, and therefore can be effectively used to alter the behavioral strategies of a network. To further explore the role of the silent neurons in the decision task, we performed a lesion experiment of the silent neurons (Fig. S5, Table S9) which revealed that lesioning of silent neurons led to more accurate networks.

## Discussion

In this work, we used a computational formalism based on RNNs to show that the natural behavioral variability observed in animals can be explained by initial configurational differences across brain networks undergoing the same training protocol. This behavioral variability reflects different strategies adopted by networks, including choice biases and different levels of speed-accuracy trade-offs^36–38^, and can be externally altered via external modulatory currents^19^. Furthermore, we showed that networks tend to choose more accurately and faster in multimodal rather than unimodal trials, highlighting the importance of multisensory integration mechanisms in perception and in strong agreement with experimental evidence and computational frameworks^10,12–15,39–41^. Adopting a multisensory decision-making task was an optimal choice for this study, due to the task incorporating not only choice but also modality properties. We would however expect to find similar variability of behavioral strategies in other complex cognitive and behavioral tasks, as long as it can be rooted on similar physiological and functional heterogeneity principles.

### The link between neural selectivity and behavioral strategies

Trained networks developed heterogeneous single-unit responses to different trial conditions, revealing interesting relationships between different selective units and adopted strategies. Fast networks had more choice- and mixed-selective units and less silent neurons than slow networks. Accurate networks, on the other hand, had significantly less choice-selective neurons compared to inaccurate ones. Combining these findings suggests that an increased number of choice-selective neurons could lead to adopting a fast-but-inaccurate strategy, whereas the number of silent and mixed-selective units play a more important role for controlling speed of different decision-making strategies without affecting its accuracy. Networks in which modality-, choice- and mixed-selective neurons are more balanced would therefore align with a slow-but-accurate behavioral strategy. Even though we observed differences in bias on an individual network level, there was not a systematic bias based on the number and type of selective units.

The association between neural selectivity and behavioral strategies was not the same for excitatory and inhibitory neurons, suggesting that both cell types play different roles in task computations. On a population level, neurons in excitatory and inhibitory populations displayed the same level of choice selectivity, in agreement with recent experimental findings^42^, and also the same level of modality selectivity^15^. We found, however, that excitatory populations generally contained a larger percentage of mixed-selective units than inhibitory populations. This might mean that excitatory neurons generally integrate more diverse information than inhibitory neurons. The difference between excitatory and inhibitory populations in hyper-selective units was small, but significant, and seemingly suggests that the inhibitory population of neurons has a larger percentage of hyper-selective units.

When considering networks with specific behavioral strategies, we observed that the high number of choice- and mixed-selective units and the low number of silent neurons in fast-decision networks (respect to slow-decision ones) could be attributed to differences in the behavioral strategy preferences of excitatory neurons. The same holds for the case of accurate and inaccurate networks, where again the significant difference in choice-selective units seemed to result from a difference in the excitatory population rather than the inhibitory population. This could imply that networks adopt different accuracy or speed strategies based on the percentage of excitatory selective units, and that inhibitory neurons support this differentiation by an internal fine-tuning which does not require a change in the total fraction of inhibitory selective units, or by regulating the stability and competition of those circuits^35,43^.

We chose a firing rate-based approach for classifying selective units in contrast to other possible options such as a receiver-operator characteristic (ROC) approach^33,35^. We chose this approach as the rate-based framework allows for classifying more types of units (e.g., silent and hyper-selective neurons) as compared to the ROC approach (Fig. S4). We also opted to consider the different selectivity categories as exclusive (i.e. a mixed-selective neuron is not counted as also a choice-selective one); this explains the lower percentage of choice selective units as compared to other studies^33,35,42^.

### Within-subject modulation of behavioral strategies

We were able to influence a network’s strategy by adding a modulatory current targeting specific groups of neurons. Signals from other brain regions, both cortical and subcortical, have been suggested to play gating or modulatory roles in cognitive functions^44–47^, albeit their specific effects on RNNs performing complex neural computations like the ones considered here have not yet been studied in depth. Targeting different selective units resulted in different behavioral adaptations, with networks becoming faster, slower, or unchanged depending on which neuron groups were targeted. All types of possible change were also observed in terms of accuracy. As has been shown in recent work^19,20^, animals tend to alternate strategies during experimental sessions, and our results indicate that this could be caused by varying levels of external input from e.g., different brain areas targeting different sub-networks of neurons. Interestingly, targeting inhibitory neurons was the only manipulation in which networks generally became slower and more accurate, whereas mostly networks become faster and less accurate upon the application of a modulatory current. Targeting choice-selective neurons was the only condition in which the network became both faster and more accurate. The modulatory current does not seem to affect the accuracy of networks when applied to mixed selective units. For silent neurons it appeared that their number, rather than their firing-rate, influenced the network’s speed since we found that (i) fast networks have significantly less silent neurons compared to slow networks, but (ii) applying a modulatory current to silent neurons does not affect reaction times. Interestingly, forcing silent neurons to decrease their activity from spontaneous levels to zero improved network accuracy, in contrast to applying the modulatory current to silent neurons, which led to more inaccurate networks. This suggests that the low firing rate of silent neurons acts as a noise-control parameter in the system. Decreasing this noise raises accuracy in decision-making, and increasing the number of silent neurons lowers decision speed.

Overall, our study suggests that the often-overlooked inter-subject behavioral variability contains useful information and reflects individual-specific strategies which may emerge from initial differences in neural networks under the same training protocol. These strategies are flexible (i.e. can be modulated by internal brain signals) and are reflected in specific neural signatures (i.e. selectivity properties). We consider that our RNN approach will allow to explore the origins and implications of cognitive and behavioral flexibility.

## Methods

All model development, training and data analysis was done using Python 3.9 and libraries including PyTorch, NumPy, matplotlib, SciPy and Sklearn.

### RNN model description

We trained 100 Recurrent neural networks (RNNs) based on earlier work^26,28^. The RNNs receive time-varying inputs and produce two outputs coding for the network’s decision (Fig. 1a). The RNNs are described by discretized equations (or, equivalently, by differential equations whose discretization) of the form

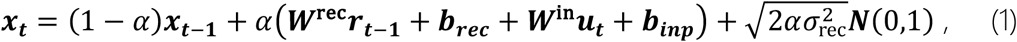

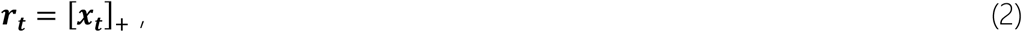

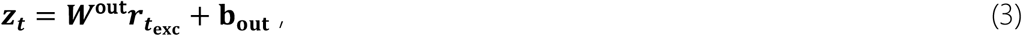

where *α* = Δ*t*/*τ* and *τ* describes the time constant of the network. In these equations, ***x***_***t***_ is a vector describing the voltage of the neurons in the network at time t, and ***r***_***t***_ describes the firing rates of the neurons at the same time, which is obtained after applying a rectified linear (ReLU) nonlinearity to the voltage, [*x*]_0_ = *max*(*x*, 0). The neuron voltage decays with a factor of *α* at each timestep and receives the firing rates of the previous timestep ***r***_***t***−**1**_ via the matrix ***W***^rec^. At every timestep, the network also receives a time-varying input ***u***_***t***_ via matrix ***W***^in^. The output variables ***z***_***t***_ are obtained through ***W***^out^, which are given randomly fixed but positive values. Note that we only read out from the firing rates of the excitatory populations as this readout can be seen as a long-range projection onto other areas of the brain, which usually involves excitatory synapses. Noise is drawn from a normal distribution ***N***(0,1) with zero mean and unit variance was drawn at every timestep and scaled accordingly. Additional constant terms ***b***_***rec***_, ***b***_***inp***_, ***b***_***out***_ describe the influence of other brain areas in the recurrent, input and output connections respectively. Each network consisted of 150 neurons of which 80% (120) was excitatory and 20% (30) was inhibitory, following Dale’s law^28^. An alternative description of our system can be given by a set of differential equations whose Euler-discretization aligns with the equations above.

The weights and biases of the input and recurrent computations were randomly initialized from a uniform distribution 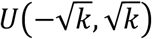. with 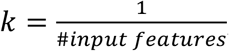, where #input features is 5 and 150 for the input and recurrent units respectively. For the output weights and biases we used *k* = 0.01. Based on trial-and-error, we multiplied the initial weights of the inhibitory units in the recurrent matrix with 6 to improve convergence.

The details of the task specific input will be discussed in the next section, but the networks received an input of the form

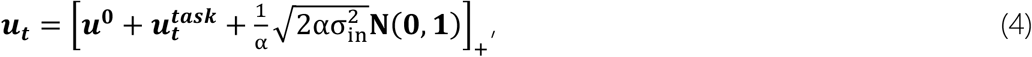

where ***u***^**0**^ represents a baseline input, 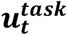 the task specific input at timestep t, and noise was drawn from a normal distribution ***N***(0,1) with zero mean and unit variance at every timestep and scaled accordingly. To ensure that the inputs are non-negative a rectified linear (ReLU) nonlinearity is applied.

### Task description

We have trained the RNNs on a multisensory integration task similar to the task described in Raposo and colleagues’ experimental work^11,33^. In this task, a rat sat in front of a monitor with a speaker which showed visual flashes and presented auditory clicks respectively (Fig. 1b). The flashes and clicks were presented at different frequencies between 9 Hz and 16 Hz, and the animal needed to decide if the frequency was higher or lower than an experimenter-imposed threshold. Rats reported their decision by moving to one of two choice ports, where one port rewarded a decision for ‘higher-than-threshold’ and the other for ‘lower-than-threshold’.

We implemented a computational version of this task^28^. The networks received inputs coding for visual and auditory frequencies ranging between 9 Hz and 16 Hz. On unimodal trials, the network just received visual or auditory information, whereas on multimodal trials the network received both visual and auditory information about the frequency. When both modalities were presented, they were congruent. The network needed to decide if the frequency presented is higher or lower than the average frequency, or 12.5 Hz. The stimulus was presented for 1 second after a fixation period of 100 ms in which no stimulus was presented. The network reported a decision by raising one of the output variables while keeping the other output variable low and could report this at any time step during the trial. We registered a decision when the difference between the two output variables was larger than 0.2. A difference of 0.2 is chosen to mark a decision because a larger value increased the number of trials in which no decision is made, whereas a threshold lower than 0.2 increased the number of trials in which a decision is already made before the onset of the stimulus (Fig. S1). We both positively- and negatively tuned the inputs in order to improve the training^28^. After the models were trained, we generated 8192 trials, where the modality and the frequency were randomly drawn with equal probability across all conditions. For each trial we saved the network’s output and firing rates at every timestep.

## RNN training

The RNNS were trained to minimize the difference between their output variables *z* and target values *T*. During the fixation period of the task, both output variables were trained to maintain a low value of 0.2. During the stimulus, the model should raise the correct output variable to a value of 1, while maintaining the other variable at the low value of 0.2. A training batch consisted of *N*_*tra****i****ls*_ = 20 trials which were randomly generated at every epoch. After the random initialization of the networks, only the weights of ***W***^rec^ and the offsets ***b***_***rec***_, ***b***_***inp***_, ***b***_***out***_ were trained to stimulate the network to use the recurrent connections, rather than the input- and output connections, to solve the task. The weights and biases were trained using back-propagation through time (BPTT) to minimize the loss function

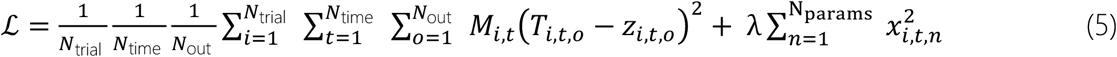

where ***M***_*i,t*_ is used as a mask for certain timepoints by using a value of 0 for timepoints that need to be excluded from the loss function and λ is the term that controls the L2-regularization to stimulate the networks to maintain small parameter values. The first 200 ms after stimulus onset were excluded from the loss function during training to avoid punishing the model for not directly making a decision and take into account biologically plausible processing times^26,28,31^. We performed the BPTT and parameter updates using stochastic gradient descent with the use of PyTorch’s SGD optimizer with a learning rate of 0.01 and MSEloss(), where the L2-regularization was implemented using the weight_decay() parameter of the SGD optimizer with a value of 0.1. In order to satisfy Dale’s law, excitatory weights that became negative or inhibitory weights that became positive during the updating step were reset to zero immediately after^28^.

The networks were trained until they reached sufficient task performance, with a maximum of 10,000 epochs. If a model did not reach sufficient task performance within 10,000 epochs, or 200,000 trials, the training was terminated and model was not used for experimentation. Sufficient task performance was defined by two conditions: (1) at least 90% of trials was valid (no decision made before stimulus onset) and (2) the network made a correct choice in at least 80% of all valid trials. To assess network’s task performance during training we generated a validation batch of *N*_*trails*_ = 1024 trials every 500 epochs and measured network performance. An overview of the training parameters is shown in Table 1.

**Table 1.**
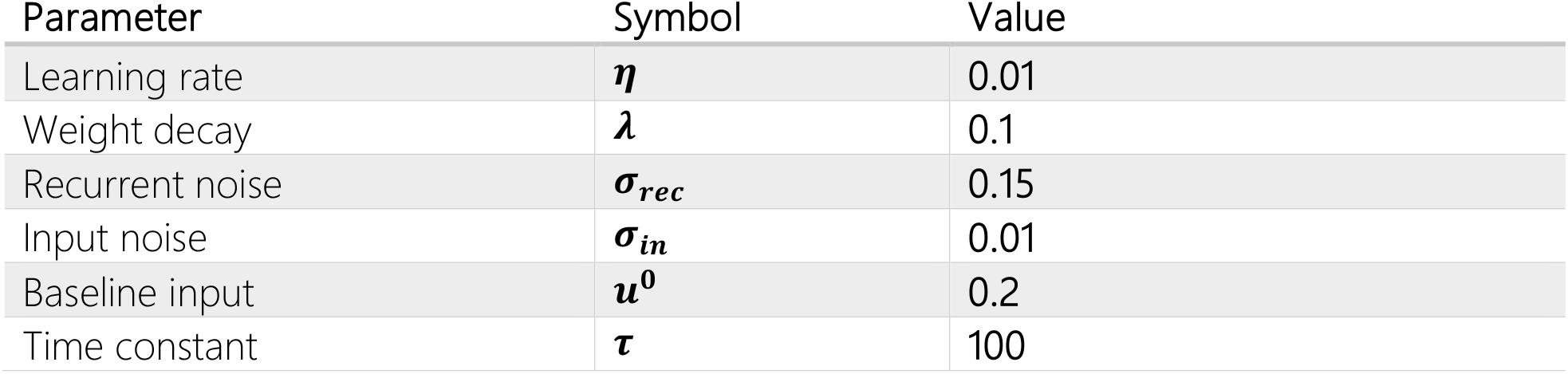
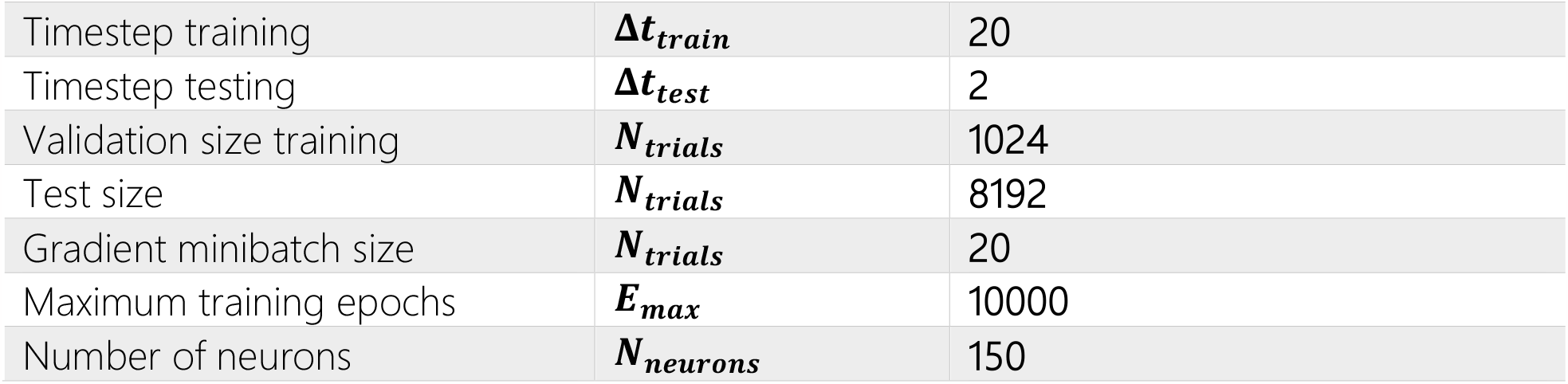
Parameter overview used for training and network design. All networks were trained with the parameters that are listed below. After random initialization, networks developed different structures following the same training procedure.

### Performance metrics and strategy mapping

For each network we computed the psychometric function based on trials in which a decision was made during the stimulus presentation. Trials in which no decision was made, or a decision was made before the stimulus onset, were excluded from the analysis. For each frequency we computed the fraction of trials in which the network reported that the frequency was higher than 12.5 Hz and fit a sigmoidal function of the form

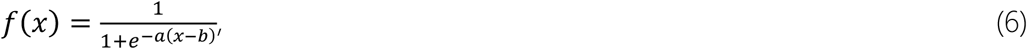

where a is a slope parameter of the sigmoid and *b* the bias, with the use of the *curve_fit()* function from SciPy. We fitted this function to the difference of the frequency and the threshold of 12.5 Hz, resulting in a range of -3.5 Hz to 3.5 Hz centered around 0. We took the a of the fitted function as a measure of the accuracy of a network, as a higher a indicates a more abrupt change in choice from ‘lower-than-threshold’ to ‘higher-than-threshold’ frequency (Fig 3a, left). We took the *b* of the fitted function as a measure of bias, as a larger *b* indicates that a model shifts its sigmoidal curve to the right, hence changes from a ‘lower-than-threshold’ to ‘higher-than-threshold’ choice at a larger frequency than the 12.5 Hz (Fig 3a, left). The sigmoidal curve was fitted both per modality and across modalities. The former is done to show how models performed on unimodal versus multimodal trials in terms of accuracy and choice bias, whereas the latter is done to assess how accurate and biased a network is in general, which is used to investigate the different strategies that models adopt.

Alternatively, we have fitted a sigmoidal function with lapse parameters of the form

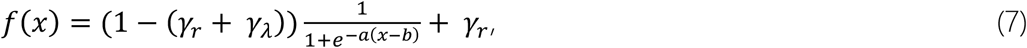

where γ_*r*_ and γ_λ_ account for guesses and lapses at the lowest and highest frequencies respectively. Lapses are defined as the fraction of ‘high’ choices at the lowest possible frequency, or the fraction of ‘low’ choices at the highest possible frequency^19,48^ (Fig. S3c).

For each network we also computed the chronometric function based on all correct trials. We calculated the average reaction time of the networks for different distances to the threshold frequency of 12.5 Hz (Fig 3a, center). We calculated this per modality to investigate the differences in reaction time between unimodal and multimodal trials. We took the average reaction time across all correct trials as a measure of how fast a network is in general, which is used to investigate the different strategies that models adopt. In order to rescale the calculated decision times to milliseconds, we multiplied our decision time, which is just the index of the time step, with Δ*t*. To account for possible processing time of sensory inputs and motor outputs that are needed for decision making in a real biological system, we added a constant value of 200 ms to the obtained reaction times in order to report more biologically plausible reaction times.

To map these behavioral metrics to strategies, we labeled a network as fast if the network has a reaction time that is less than the average reaction time of all networks, and slow otherwise. We labeled a network as accurate if the slope of the fitted sigmoidal curve is larger than the average slope of all networks, and inaccurate otherwise.

### Measuring unit selectivity

Based on the firing rates of the units on correct trials^33^ we assessed the selectivity of the neurons. Before the neurons were classified, the average firing rate across trial conditions was calculated. The trial conditions of interest were ‘auditory-high’, ‘auditory-low’, ‘visual-high’ and ‘visual-low’. The multimodal trials were excluded for this analysis as it would be ambiguous to attribute selectivity when two conditions were presented simultaneously. We defined five types of selectivities (Fig. 4a):

- **Modality**: when a neuron fires most on either auditory or visual trials irrespective of the frequency of the trial. For example, when a neuron fires more on auditory trials with a high and low frequency as compared to visual trials with either frequency, it is classified as modality-selective.
- **Choice**: when a neuron fires more for high- or low-frequency trials as compared to the opposite. For example, when a neuron fires more for auditory- and visual trials with a high frequency compared to auditory- and visual trials with a low frequency. Note that this is identical to firing more for a specific choice (high or low) since we only looked at correct trials.
- **Mixed**: when a neuron is both modality- or choice-selective. We do this by first classifying the neuron into one type (modality or choice) and then testing for the other one. In case of an initially-classified-as modality-selective neuron, we define this by looking if the firing rates also show separation based on high-versus low-frequency trials. This is done by taking the average of the firing rates for the high-frequency trials and separately the average of the firing rates for the low-frequency trials and measuring if the difference between those two is larger than the difference between the 2^nd^ and 3^rd^ largest firing rate conditions. In case of an initially-classified-as choice-selective neuron, this procedure is similar but the difference between the average firing rates of visual versus the average firing rates of auditory trials is compared against the difference between the 2^nd^ and 3^rd^ largest firing rate conditions.
- **Silent**: when a neuron fires less during the stimulus than the maximum firing rate during the fixation period for all unimodal trial types.
- **Hyper-selectivity**: when a neuron fires less during the stimulus than the maximum firing rate during the fixation period for three out of four unimodal trial types, but fires more for one, e.g., auditory high-frequency trials only.

We used the average firing rate over the last 20 timepoints, or 40 ms, of the trials to assess a neuron’s selectivity.

Alternatively, we computed the receiver operating characteristic^33,35^ (ROC) and the area under this curve (AUC_ROC_) using the using roc_curve() and auc() functions from Sklearn. The same timepoints as in our original approach were used to determine the selectivities of the neurons. A neuron with no selectivity for modality or choice would have an AUC_ROC_ of 0.5. We determined if a neuron was selective by comparing the AUC_ROC_ of the neuron against the AUC_ROC_ of a shuffled distribution and marked a neuron selective if the AUC_ROC_ was in the lowest or highest 2.5 percentile of this shuffled distribution. In case of choice selectivity, the AUC_ROC_ was calculated based on the network’s choice, and for the shuffled distribution the choices were shuffled for 1000 times, while keeping the network’s activity the same. In case of modality selectivity, the trials were first split in trials ending in leftward and rightward choices. Then for each group, we checked how well the neuron’s activity predicts the trial modality (if it was visual or auditory). If in either the leftward or rightward trials the AUC_ROC_ differed significantly from a shuffled distribution in which the trial modalities are randomly shuffled, the neuron is marked as modality selective. Neurons that are marked both choice- and modality selective are labeled mixed selective.

### Adding modulatory current to selective units

To investigate to what extent we could change a network’s strategy, we added a modulatory current to specific neurons of each network during the trial, which experimentally could be achieved using optogenetics. In total 8 different experiments were performed targeting different sets of neurons. The 8 sets that were targeted were (1) modality selective neurons, (2) choice selective neurons, (3) mixed selective neurons, (4) silent neurons, (5) hyper-selective neurons, (6) all neurons, (7) inhibitory neurons, (8) excitatory neurons. We modulated the system by adding a constant bias current with noise to the targeted neurons, changing the equation of the current of the targeted neurons to

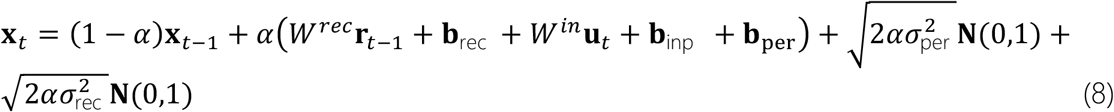

where a constant bias ***b***_*per*_ of 0.2 is added and random noise sampled at every timestep with σ_*per*_ = 0.01. We generated *N*_*trails*_ = 2048 trials per modulatory setup and added a baseline experiment of the same number of trials with no modulatory current against which the results of the modulatory currents were compared in terms of psychometric- and chronometric function. We added the modulatory current throughout the entire trial duration.

### Lesion experiment

To investigate the role of silent neurons in the network, we performed an additional experiment where we effectively lesioned silent neurons from the network. Our definition of silent neurons allows for the neurons that are marked silent to not be completely silent but rather have a firing rate lower than in the fixation period. To see if these neurons have any function, we perform an experiment in which we set the firing rates of those silent neurons to exactly zero, effectively lesioning them from the network. We compare the results of those lesioned networks against their baseline performance in terms of psychometric and chronometric function by generating *N*_*trails*_ = 2048 trials.

### Statistical analyses

We used permutation tests of the difference in means with n=100,000 resamples to assess if there were significant differences between groups (i.e. modality, choice, mixed, excitatory, inhibitory, etc). In case of comparisons between groups assigned to different strategies, we used the independent permutation test of SciPy which tests the null hypothesis that all observations are sampled from the same underlying distribution. For the experiments in which we apply a modulatory current to the networks, we used the paired samples alternative of the permutation test, which has the null hypothesis that the observations within each pair representing the same network before and after modulation is drawn from the same underlying distribution. To correct for the multiple comparisons that we made, we applied the Holm-Bonferroni correction. For the comparisons made between strategies (Fig.4) and the total inhibitory versus excitatory population (Fig. 5a) we corrected for the 5 comparisons made on the same groups, whereas for the excitatory vs inhibitory comparisons (Fig. 5b and c) we corrected for the 20 comparisons made. For the correlation between the number of selective units and the different metrics of behavior, we calculated the Spearman correlation coefficient with associated p-value. Raw and adjusted p-values can be found in the tables available in the Supplementary material.

## Acknowledgements

The authors thank Amparo Gilhuis and Guangyu Yang for insightful discussions in early stages of this project.

## Author contributions

JFM conceived and designed the study; TW and SD performed the research; TW, SD and JFM analyzed the results; TW, SD, CP and JFM wrote the manuscript.

## Data and materials availability

All information needed to reproduce the results of this manuscript are in the main text and Methods section. The code used to generate the results will be made available upon publication of this work.

## Supplementary material

### Supplementary figures

**Figure S1.**
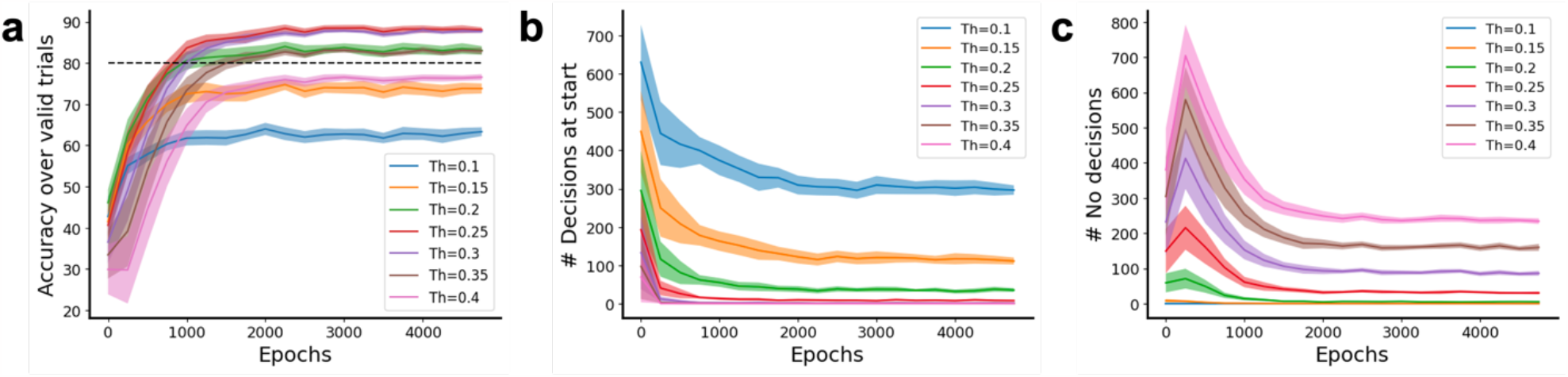
Decision threshold influence on classification accuracy and valid trials. **a** We explored the effect of different thresholds for the difference between the output variables that is used to mark a network’s decision time. We observed that the lower the threshold, the more errors the networks make and that the classification accuracy increases until a threshold of 0.3, after which it drops again, likely due to less decisions being made at higher thresholds. The candidate threshold range that achieved a classification accuracy over 80% were 0.2 – 0.35. **b** The lower the imposed threshold, the more decisions the model already made before the stimulus onset, which we marked as invalid trials. Especially thresholds below 0.2 result in a relatively large fraction of trials becoming invalid. **c** The number of trials in which no decision is made, so where the output variables of the networks do not separate enough, increases with the decision threshold. Especially thresholds larger than 0.25 result in a relatively large fraction of trials becoming invalid.

**Figure S2.**
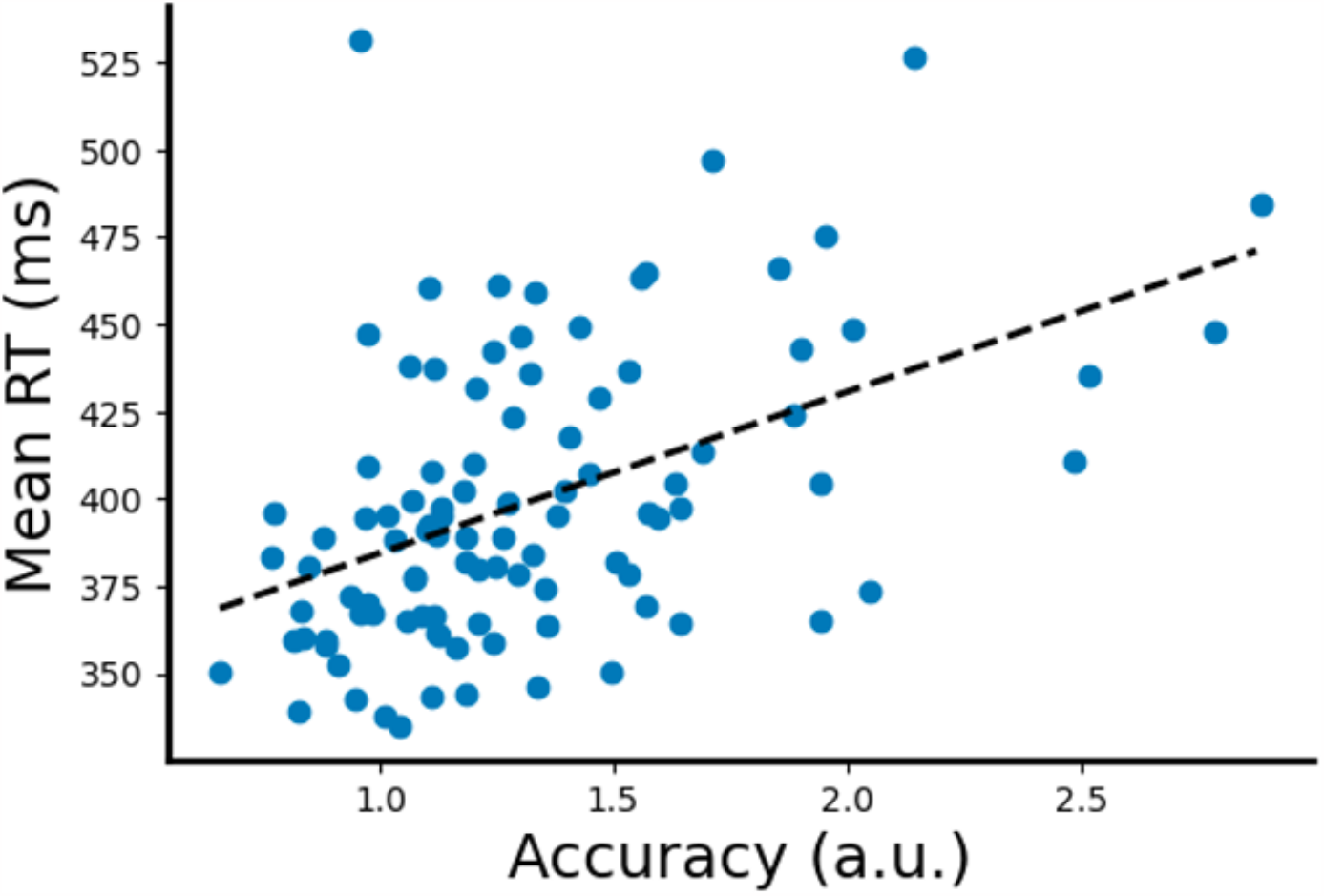
Speed-accuracy trade-off. A positive correlation of 0.47 (p < 0.001, Pearson R) is apparent between the accuracy of networks and the mean reaction time, in line with earlier reports of a speed-accuracy trade-off reported in both animal and computational studies.

**Figure S3.**
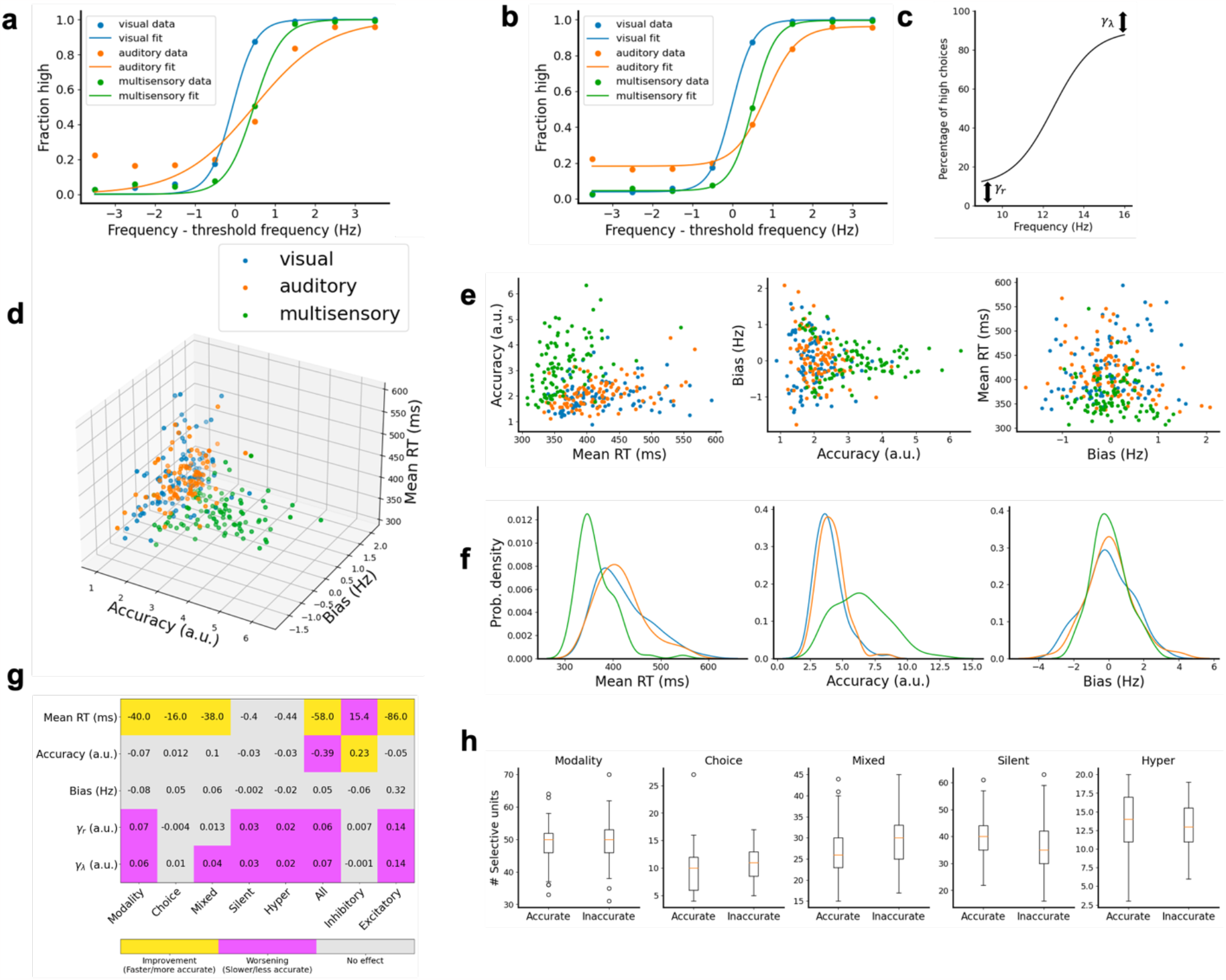
Summary of results when psychometric functions are fitted with lapse parameters. **a** Example of a standard sigmoidal function fitted to data of a network as used in this research. **b** Example of the same data as in **a** but fitted with a sigmoidal function with lapse parameters. **c** Lapse parameters indicate the number of high-choices at the lowest frequency (γ_λ_) and low-choices at the highest frequency (γ_r_), changing the range of the psychometric function from (0, 1) to (γ_λ_, 1 − γ_λ_). **d** Similarly to Fig. 3b, we observe spread in all metrics of behavior where high accuracy and low mean reaction times is mostly achieved on multisensory trials. **e** Similar to Fig. 3c, 2D projections of **d** reveal that networks achieve low reaction times on multisensory trials (left) while still achieving high accuracy. The bias also seems to be reduced with a better accuracy (center) which is mostly achieved on multisensory trials. The bias level does not seem to change a lot with reaction times, but it is again apparent that most low reaction times are achieved on multisensory trials. **f** Distribution plots of **e** show again an average lower reaction time on multisensory trials as compared to unimodal trials (left), a wider spread in accuracy on multisensory trials (center) and a narrower distribution of bias levels on multisensory trials (right). The difference between multisensory and unimodal bias seems smaller compared to Fig. 3d. **g** Fitting psychometric curves to the data obtained after application of a modulatory current reveals that often there is no significant effect on the accuracy (slope) directly, but that often there is a shift in lapse parameters that can account for the observed change in accuracy in the standard sigmoidal fitting. See Table S1. **h** When looking at the difference between accurate and inaccurate networks based on the accuracy of networks after fitting with a sigmoidal function with lapse parameters, we see no significant changes in choice selective neurons anymore, as observed in Fig.4c. However, the slope of this sigmoid might not capture the accuracy as explicit anymore since the lapse parameters are not included in the definition of accuracy in this situation and a more elaborate definition of accuracy would be better suitable.

**Figure S4.**
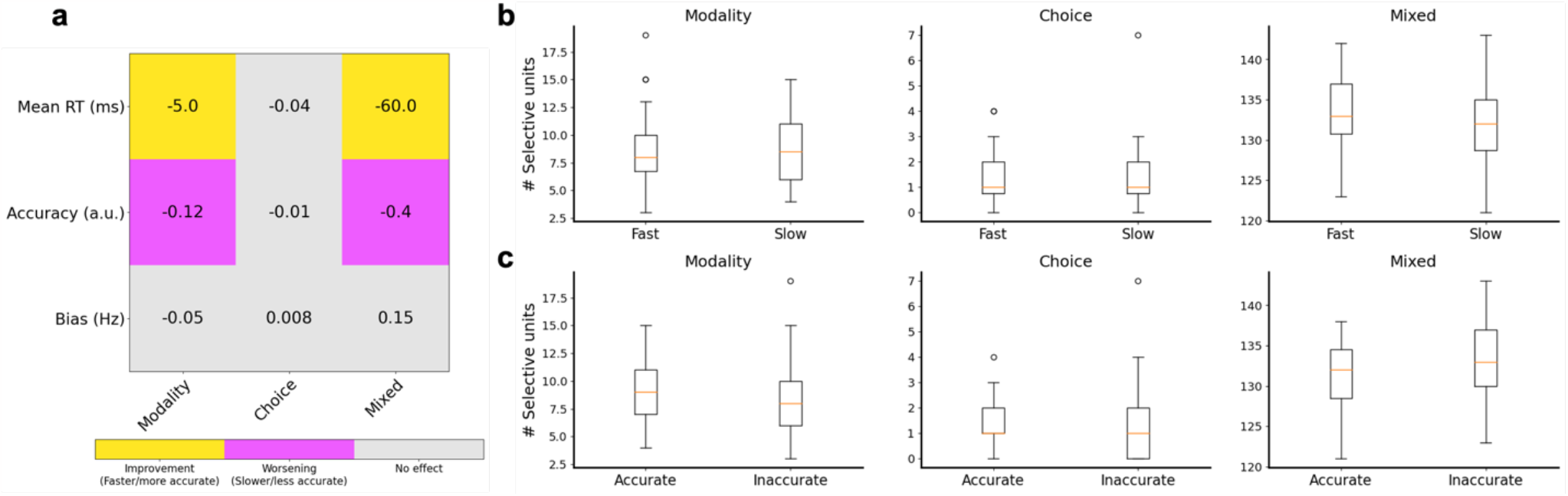
ROC analysis to mark selectivity results in mostly mixed-selective units. **a** Adding a modulatory current to selective units based on the ROC method results in no significant effects for choice selective units. The reason for this is likely because of the low number of neurons that are marked as pure-choice selective, as most neurons are marked to be mixed selective following the ROC approach. The large number of mixed selective units also results in a significant decrease in accuracy when these neurons are targeted as compared to Fig. 6c where there was no significant effect visible after targeting mixed selective units, likely because there were less mixed selective units present. **b** There do not seem to be any significant differences between fast and slow groups using the ROC definition, likely because almost all neurons are marked to be mixed selective as compared to our rate-based classification approach. **C** Same as **b** but for accurate versus inaccurate networks.

**Figure S5.**
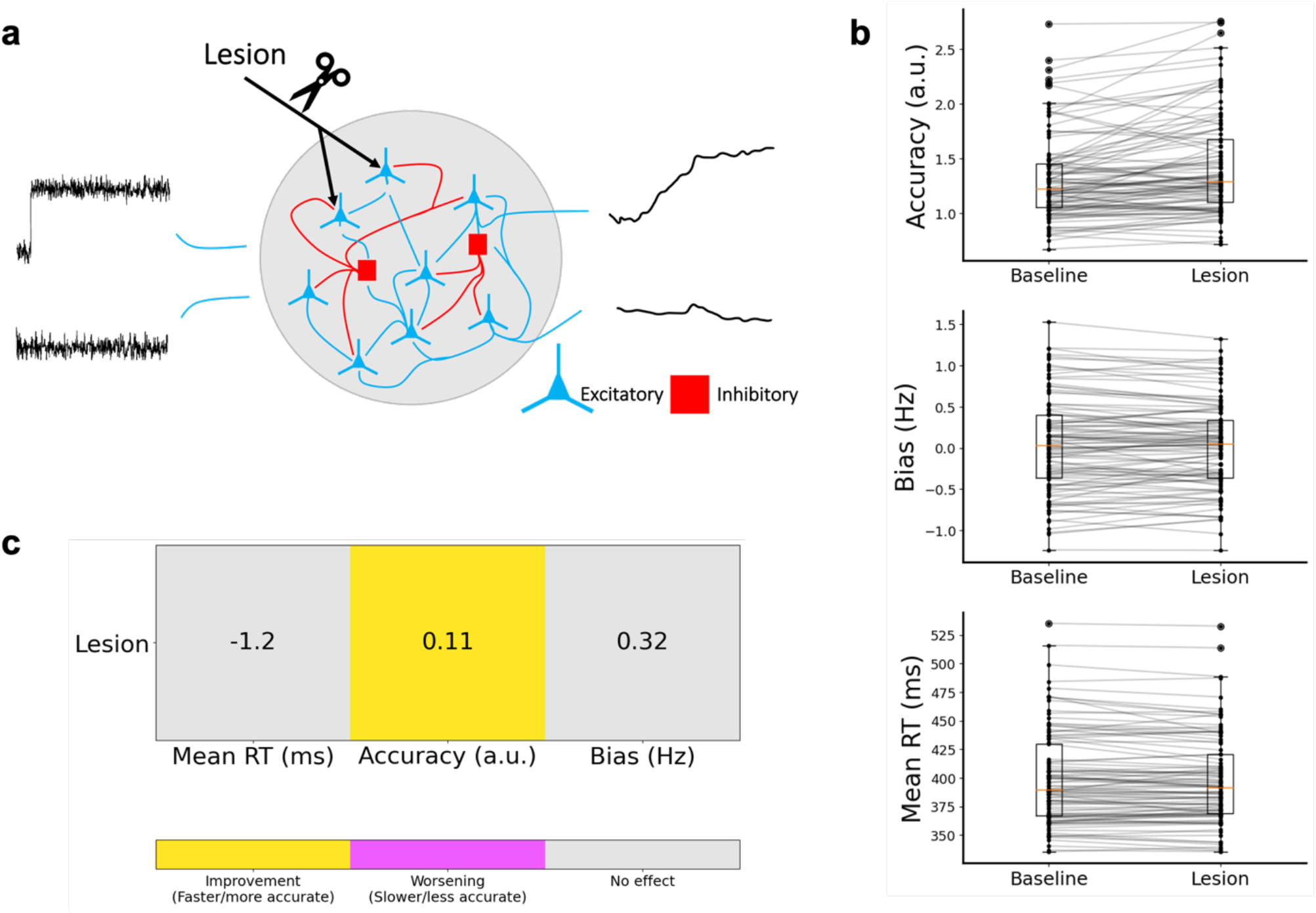
Lesioning silent neurons reveals functional role. **a** We lesioned neurons that were silent by forcing their firing rate to be zero at every timestep. **b** Comparing the networks’ performances before and after lesioning reveals that only the accuracy is significantly increased by lesioning the silent neurons. **c** Lesion of the silent neurons improves accuracy but does not significantly affect the mean reaction time or the bias.

### Supplementary Tables

**Table S1:** Raw p-values based on the permutation test for the application of a modulatory current as compared to baseline performance with lapse fitting (Fig. S3g).

| | *Slope* | *Offset* | *Mean RT* | $\gamma_r$ | $\gamma_\lambda$ |
| --- | --- | --- | --- | --- | --- |
| *Modality* | 0.803871961 | 0.35097649 | 1.99998E-05 | 1.99998E-05 | 1.99998E-05 |
| *Choice* | 0.783672163 | 0.544334557 | 1.99998E-05 | 0.644593554 | 0.288117119 |
| *Mixed* | 0.865191348 | 0.618733813 | 1.99998E-05 | 0.281637184 | 0.008779912 |
| *Silent* | 0.400435996 | 0.925330747 | 0.638213618 | 1.99998E-05 | 1.99998E-05 |
| *Hyper* | 0.445255547 | 0.30501695 | 0.60601394 | 0.000159998 | 1.99998E-05 |
| *All* | 1.99998E-05 | 0.696353036 | 1.99998E-05 | 0.000259997 | 1.99998E-05 |
| *Inhibitory* | 1.99998E-05 | 0.430095699 | 1.99998E-05 | 0.246197538 | 0.876691233 |
| *Excitatory* | 0.909610904 | 0.364396356 | 1.99998E-05 | 1.99998E-05 | 1.99998E-05 |

**Table S2:** Raw and adjusted p-values based on the permutation test and Holm-Bonferroni correction for difference in strategies (Fig 4b and c).

|  | *Fast versus slow* |  | *Accurate versus inaccurate* |  |
| --- | --- | --- | --- | --- |
|  | *p-value* | *Adjusted p-value* | *p-value* | *Adjusted p-value* |
| *Modality* | 0.0666593 | 6.66593334e-02 | 0.530794 | 1 |
| *Choice* | 0.0095199 | 2.85597144e-02 | 0.001179988 | 0.00589994 |
| *Mixed* | 1.99998e-05 | 9.99990000e-05 | 0.054539 | 0.16361836 |
| *Silent* | 3.99998e-05 | 1.59999200e-04 | 0.0349996 | 0.1399986 |
| *Hyper* | 0.03253967 | 6.50793492e-02 | 0.974470 | 1 |

**Table S3:** Correlations between the number of selective units and metrics of behavior (Spearman’s rho).

|  | *Slope* | *Offset* | *Mean RT* |
| --- | --- | --- | --- |
| *Modality* | -0.05 (p: 0.6) | -0.063 (p: 0.53) | -0.21 (p: 0.034) |
| *Choice* | -0.3 (p: < 0.01) | 0.1 (p: 0.32) | -0.42 (p: < 0.001) |
| *Mixed* | -0.29 (p: < 0.01) | 0.038 (p: 0.7) | -0.45 (p: < 0.001) |
| *Silent* | 0.36 (p: < 0.001) | -0.07(p: 0.49) | 0.49 (p: < 0.001) |
| *Hyper* | 0.18 (p: 0.08) | -0.026 (p: 0.80) | 0.32 (p: < 0.01) |

**Table S4:** Raw and adjusted p-values based on the permutation test and Holm-Bonferroni correction for difference in EI (Fig. 5a).

|  | Inhibitory versus Excitatory |  |
| --- | --- | --- |
|  | p-value | Adjusted p-value |
| Modality | 8.85951140e-01 | 8.85951140e-01 |
| Choice | 3.12996870e-02 | 9.38990610e-02 |
| Mixed | 5.99994000e-05 | 2.99997000e-04 |
| Silent | 9.61590384e-02 | 1.92318077e-01 |
| Hyper | 7.73992260e-03 | 3.09596904e-02 |

**Table S5:** Raw and adjusted p-values based on the permutation test and Holm-Bonferroni correction for difference in strategies between EI related to speed (Fig. 5b).

|  | Inh. Fast v Exc. Fast |  | Inh. Fast v Inh. Slow |  |
| --- | --- | --- | --- | --- |
|  | p-value | Adjusted p-value | p-value | Adjusted p-value |
| Modality | 0.320077 | 1 | 0.036700 | 0.484675 |
| Choice | 0.000280 | 0.484675 | 0.631114 | 1 |
| Mixed | 0.000060 | 0.001200 | 0.227578 | 1 |
| Silent | 0.021420 | 0.321297 | 0.087859 | 1 |
| Hyper | 0.087759 | 1 | 0.105699 | 1 |

|  | Exc. Fast v Exc. Slow |  | Exc. Slow v Inh. Slow |  |
| --- | --- | --- | --- | --- |
|  | p-value | Adjusted p-value | p-value | Adjusted p-value |
| Modality | 0.280737 | 1 | 0.337277 | 1 |
| Choice | 0.000280 | 0.484675 | 0.920411 | 1 |
| Mixed | 0.000100 | 0.001800 | 0.100279 | 1 |
| Silent | 0.000060 | 0.001200 | 0.978070 | 1 |
| Hyper | 0.123819 | 1 | 0.034620 | 0.484675 |

**Table S6:** Raw and adjusted p-values based on the permutation test and Holm-Bonferroni correction for difference in strategies between EI related to accuracy (Fig 5c).

|  | Inh. Accurate v Exc. Accurate |  | Inh. Accurate v Inh. Inaccurate |  |
| --- | --- | --- | --- | --- |
|  | p-value | Adjusted p-value | p-value | Adjusted p-value |
| Modality | 0.881871 | 1 | 1.000000 | 1 |
| Choice | 0.371556 | 1 | 0.405376 | 1 |
| Mixed | 0.072499 | 0.956190 | 1.000000 | 1 |
| Silent | 0.685693 | 1 | 0.482975 | 1 |
| Hyper | 0.175338 | 1 | 0.800892 | 1 |

|  | Exc. Accurate v Exc. Inaccurate |  | Exc. Inaccurate v Inh. Inaccurate |  |
| --- | --- | --- | --- | --- |
|  | p-value | Adjusted p-value | p-value | Adjusted p-value |
| Modality | 0.467095 | 1 | 0.746493 | 1 |
| Choice | 0.000440 | 0.008360 | 0.041240 | 0.659833 |
| Mixed | 0.044080 | 0.661193 | 0.000380 | 0.007600 |
| Silent | 0.032360 | 0.550114 | 0.068299 | 0.956190 |
| Hyper | 0.859791 | 1 | 0.021080 | 0.379436 |

**Table S7:**
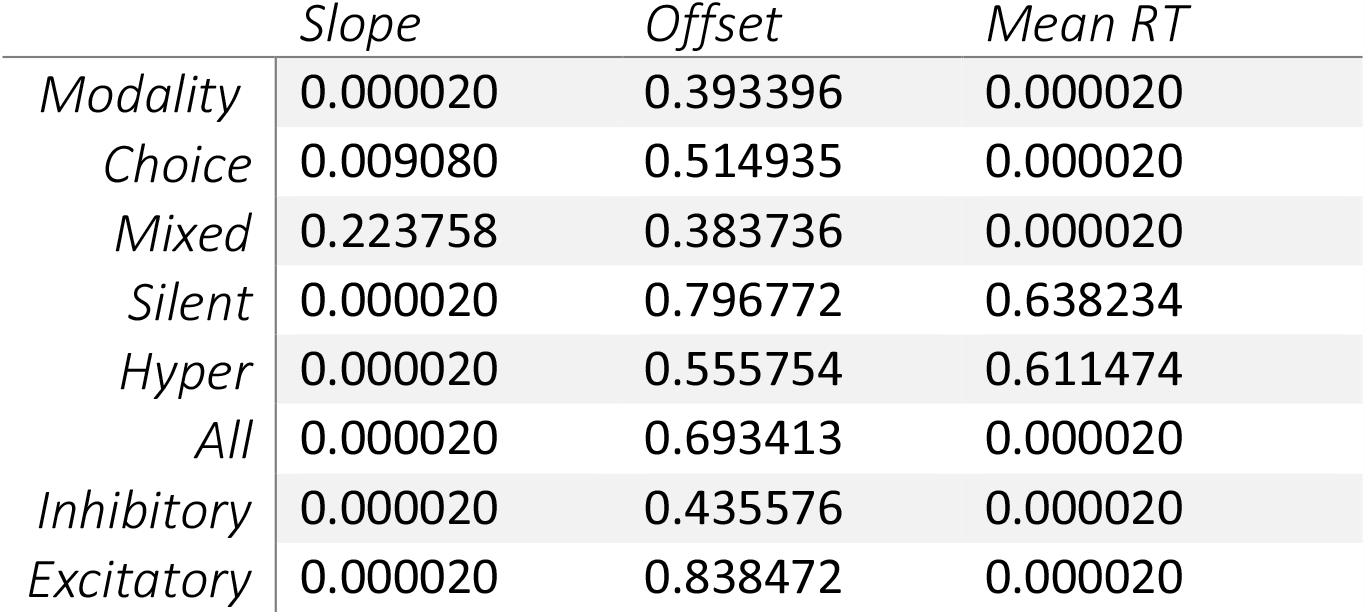
Raw p-values based on the permutation test for the application of a modulatory current as compared to baseline performance (Fig. 6c).

|  | Slope | Offset | Mean RT |
| --- | --- | --- | --- |
| Modality | 0.000020 | 0.393396 | 0.000020 |
| Choice | 0.009080 | 0.514935 | 0.000020 |
| Mixed | 0.223758 | 0.383736 | 0.000020 |
| Silent | 0.000020 | 0.796772 | 0.638234 |
| Hyper | 0.000020 | 0.555754 | 0.611474 |
| All | 0.000020 | 0.693413 | 0.000020 |
| Inhibitory | 0.000020 | 0.435576 | 0.000020 |
| Excitatory | 0.000020 | 0.838472 | 0.000020 |

**Table S8:**
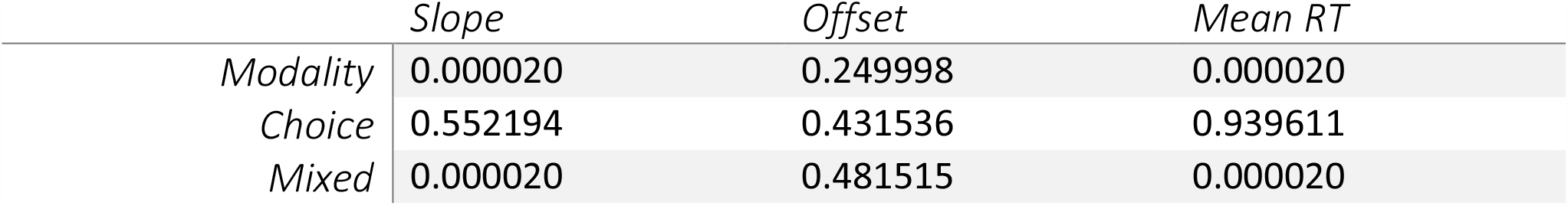
Raw p-values based on the permutation test for the application of a modulatory current as compared to baseline performance using the ROC definition (Fig. S4a).

|  | Slope | Offset | Mean RT |
| --- | --- | --- | --- |
| Modality | 0.000020 | 0.249998 | 0.000020 |
| Choice | 0.552194 | 0.431536 | 0.939611 |
| Mixed | 0.000020 | 0.481515 | 0.000020 |

**Table S9:** Raw p-values based on the permutation test for lesioning as compared to baseline performance (Fig.S 5c).

|  | *Slope* | *Offset* | *Mean RT* |
| --- | --- | --- | --- |
| *Lesion* | 0.00002 | 0.228218 | 0.092959 |

## References

1. Kanai, R. & Rees, G. The structural basis of inter-individual differences in human behaviour and cognition. Nat Rev Neurosci 12, 231–242 (2011).

2. Faure, P., Fayad, S. L., Solié, C. & Reynolds, L. M. Social Determinants of Inter-Individual Variability and Vulnerability: The Role of Dopamine. Front Behav Neurosci 16, 836343 (2022).

3. Buchanan, S. M., Kain, J. S. & de Bivort, B. L. Neuronal control of locomotor handedness in Drosophila. Proceedings of the National Academy of Sciences 112, 6700–6705 (2015).

4. Stern, S., Kirst, C. & Bargmann, C. I. Neuromodulatory Control of Long-Term Behavioral Patterns and Individuality across Development. Cell 171, 1649–1662.e10 (2017).

5. Tuttle, A. H., Philip, V. M., Chesler, E. J. & Mogil, J. S. Comparing phenotypic variation between inbred and outbred mice. Nat Methods 15, 994–996 (2018).

6. Brunton, B. W., Botvinick, M. M. & Brody, C. D. Rats and Humans Can Optimally Accumulate Evidence for Decision-Making. Science 340, 95–98 (2013).

7. Shadlen, M. N. & Newsome, W. T. Neural basis of a perceptual decision in the parietal cortex (area LIP) of the rhesus monkey. J. Neurophysiol. 86, 1916–1936 (2001).

8. The International Brain Laboratory et al. Standardized and reproducible measurement of decision-making in mice. eLife 10, e63711 (2021).

9. Wang, X.-J. Probabilistic Decision Making by Slow Reverberation in Cortical Circuits. Neuron 36, 955–968 (2002).

10. Sheppard, J. P., Raposo, D. & Churchland, A. K. Dynamic weighting of multisensory stimuli shapes decision-making in rats and humans. Journal of Vision 13, 4–4 (2013).

11. Raposo, D., Sheppard, J. P., Schrater, P. R. & Churchland, A. K. Multisensory Decision-Making in Rats and Humans. J. Neurosci. 32, 3726–3735 (2012).

12. Chandrasekaran, C. Computational principles and models of multisensory integration. Current Opinion in Neurobiology 43, 25–34 (2017).

13. Angelaki, D. E., Gu, Y. & DeAngelis, G. C. Multisensory integration: psychophysics, neurophysiology, and computation. Current Opinion in Neurobiology 19, 452–458 (2009).

14. Meijer, G. T. et al. Neural Correlates of Multisensory Detection Behavior: Comparison of Primary and Higher-Order Visual Cortex. Cell Reports 31, 107636 (2020).

15. Oude Lohuis, M. N. et al. Multisensory task demands temporally extend the causal requirement for visual cortex in perception. Nat Commun 13, 2864 (2022).

16. Bechara, A., Dolan, S. & Hindes, A. Decision-making and addiction (part II): myopia for the future or hypersensitivity to reward? Neuropsychologia 40, 1690–1705 (2002).

17. Pittaras, E., Hamelin, H. & Granon, S. Inter-Individual Differences in Cognitive Tasks: Focusing on the Shaping of Decision-Making Strategies. Frontiers in Behavioral Neuroscience 16, (2022).

18. Pagan, M. et al. A new theoretical framework jointly explains behavioral and neural variability across subjects performing flexible decision-making. 2022.11.28.518207 Preprint at 10.1101/2022.11.28.518207 (2022).

19. Ashwood, Z. C. et al. Mice alternate between discrete strategies during perceptual decisionmaking. Nat Neurosci 25, 201–212 (2022).

20. Le, N. M. et al. Mixtures of strategies underlie rodent behavior during reversal learning. PLOS Computational Biology 19, e1011430 (2023).

21. Marsat, G. & Maler, L. Neural Heterogeneity and Efficient Population Codes for Communication Signals. Journal of Neurophysiology 104, 2543–2555 (2010).

22. Mejias, J. F. & Longtin, A. Optimal Heterogeneity for Coding in Spiking Neural Networks. Phys.Rev. Lett. 108, 228102 (2012).

23. Mejias, J. F. & Longtin, A. Differential effects of excitatory and inhibitory heterogeneity on the gain and asynchronous state of sparse cortical networks. Frontiers in Computational Neuroscience 8, (2014).

24. Zeldenrust, F., Gutkin, B. & Denéve, S. Efficient and robust coding in heterogeneous recurrent networks. PLOS Computational Biology 17, e1008673 (2021).

25. Perez-Nieves, N., Leung, V. C. H., Dragotti, P. L. & Goodman, D. F. M. Neural heterogeneity promotes robust learning. Nat Commun 12, 5791 (2021).

26. Mante, V., Sussillo, D., Shenoy, K. V. & Newsome, W. T. Context-dependent computation by recurrent dynamics in prefrontal cortex. Nature 503, 78–84 (2013).

27. Sussillo, D. & Barak, O. Opening the Black Box: Low-Dimensional Dynamics in High-Dimensional Recurrent Neural Networks. Neural Computation 25, 626–649 (2012).

28. Song, H. F., Yang, G. R. & Wang, X.-J. Training Excitatory-Inhibitory Recurrent Neural Networks for Cognitive Tasks: A Simple and Flexible Framework. PLOS Computational Biology 12, e1004792 (2016).

29. Yang, G. R., Joglekar, M. R., Song, H. F., Newsome, W. T. & Wang, X.-J. Task representations in neural networks trained to perform many cognitive tasks. Nature Neuroscience 22, 297–306 (2019).

30. Yang, G. R. & Wang, X.-J. Artificial Neural Networks for Neuroscientists: A Primer. Neuron 107, 1048–1070 (2020).

31. Kleinman, M., Chandrasekaran, C. & Kao, J. A mechanistic multi-area recurrent network model of decision-making. in Advances in Neural Information Processing Systems (eds. Ranzato, M., Beygelzimer, A., Dauphin, Y., Liang, P. S. & Vaughan, J. W.) vol. 34 23152–23165 (Curran Associates, Inc., 2021).

32. Yang, G. R. & Mazon, M. M. Next-generation of recurrent neural network models for cognition. Preprint at 10.31234/osf.io/w34n2 (2021).

33. Raposo, D., Kaufman, M. T. & Churchland, A. K. A category-free neural population supports evolving demands during decision-making. Nat Neurosci 17, 1784–1792 (2014).

34. Rigotti, M. et al. The importance of mixed selectivity in complex cognitive tasks. Nature 497, 585– 590 (2013).

35. Roach, J. P., Churchland, A. K. & Engel, T. A. Choice selective inhibition drives stability and competition in decision circuits. Nat Commun 14, 147 (2023).

36. Heitz, R. P. & Schall, J. D. Neural Mechanisms of Speed-Accuracy Tradeoff. Neuron 76, 616–628 (2012).

37. Heitz, R. P. The speed-accuracy tradeoff: history, physiology, methodology, and behavior. Frontiers in Neuroscience 8, (2014).

38. Standage, D., Blohm, G. & Dorris, M. C. On the neural implementation of the speed-accuracy trade-off. Frontiers in Neuroscience 8, (2014).

39. Alais, D., Newell, F. N. & Mamassian, P. Multisensory processing in review: from physiology to behaviour. Seeing Perceiving 23, 3–38 (2010).

40. Meijer, G. T., Pie, J. L., Dolman, T. L., Pennartz, C. M. A. & Lansink, C. S. Audiovisual Integration Enhances Stimulus Detection Performance in Mice. Frontiers in Behavioral Neuroscience 12, (2018).

41. Meijer, G. T., Mertens, P. E. C., Pennartz, C. M. A., Olcese, U. & Lansink, C. S. The circuit architecture of cortical multisensory processing: Distinct functions jointly operating within a common anatomical network. Progress in Neurobiology 174, 1–15 (2019).

42. Najafi, F. et al. Excitatory and Inhibitory Subnetworks Are Equally Selective during Decision-Making and Emerge Simultaneously during Learning. Neuron 105, 165–179.e8 (2020).

43. Wong, K.-F. & Wang, X.-J. A recurrent network mechanism of time integration in perceptual decisions. J. Neurosci. 26, 1314–1328 (2006).

44. Murray, J. D., Jaramillo, J. & Wang, X.-J. Working Memory and Decision-Making in a Frontoparietal Circuit Model. J. Neurosci. 37, 12167–12186 (2017).

45. Mejias, J. F. & Wang, X.-J. Mechanisms of distributed working memory in a large-scale network of macaque neocortex. eLife 11, e72136 (2022).

46. Jaramillo, J., Mejias, J. F. & Wang, X.-J. Engagement of Pulvino-cortical Feedforward and Feedback Pathways in Cognitive Computations. Neuron 101, 321–336.e9 (2019).

47. Lindeman, S. et al. Cerebellar Purkinje cells can differentially modulate coherence between sensory and motor cortex depending on region and behavior. PNAS 118, (2021).

48. Prins, N. The psychometric function: The lapse rate revisited. Journal of Vision 12, 25 (2012).

